# Multimodal Data Integration Advances Longitudinal Prediction of the Naturalistic Course of Depression and Reveals a Multimodal Signature of Disease Chronicity

**DOI:** 10.1101/2023.01.10.523383

**Authors:** Philippe C. Habets, Rajat M Thomas, Yuri Milaneschi, Rick Jansen, Rene Pool, Wouter J Peyrot, Brenda WJH Penninx, Onno C Meijer, Guido A van Wingen, Christiaan H. Vinkers

## Abstract

The ability to individually predict disease course of major depressive disorder (MDD) is essential for optimal treatment planning. Here, we use a data-driven machine learning approach to assess the predictive value of different sets of biological data (whole-blood proteomics, lipid-metabolomics, transcriptomics, genetics), both separately and added to clinical baseline variables, for the longitudinal prediction of 2-year MDD chronicity (defined as presence of MDD diagnosis after 2 years) at the individual subject level. Prediction models were trained and cross-validated in a sample of 643 patients with current MDD (2-year chronicity n = 318) and subsequently tested for performance in 161 MDD individuals (2-year chronicity n = 79). Proteomics data showed best unimodal data predictions (AUROC = 0.68). Adding proteomic to clinical data at baseline significantly improved 2-year MDD chronicity predictions (AUROC = 0.63 vs AUROC = 0.78, p = 0.013), while the addition of other -omics data to clinical data did not yield significantly increased model performance. SHAP and enrichment analysis revealed proteomic analytes involved in inflammatory response and lipid metabolism, with fibrinogen levels showing the highest variable importance, followed by symptom severity. Machine learning models outperformed psychiatrists’ ability to predict two-year chronicity (balanced accuracy = 71% vs 55%). This study showed the added predictive value of combining proteomic, but not other -omic data, with clinical data. Adding other -omic data to proteomics did not further improve predictions. Our results reveal a novel multimodal signature of MDD chronicity that shows clinical potential for individual MDD disease course predictions from baseline measurements.

## Introduction

Major Depressive Disorder (MDD) is a heterogenous disorder where both treatment response and prognosis vastly differ between individuals. Around 20–25% of MDD patients are at risk for chronic depression, independent of initial treatment type (1). The ability to individually predict disease course early on is essential for optimal treatment planning, as this could allow for early treatment intensification for patients with a low long-term chance of remission, and potentially bypassing initial first-choice treatments.

Previous studies have yielded insights in clinical, psychological and biological markers for chronicity in depression. Chronicity in depression has been related to longer symptom duration, increased symptom severity and earlier age of onset (1,2); higher levels of neuroticism and lower levels of extraversion and conscientiousness (3); and various inflammatory markers (4), low levels of vitamin D (5), metabolic syndrome (6) and lower cortisol awakening response (7). Yet, statistically significant differences on a group level will not always be useful for prediction of disease course for the individual, either due to low effect sizes or redundancy with respect to other more predictive variables. While multiple studies showed biological data can be used to make accurate diagnostic predictions of MDD cases and healthy controls (8–10), individual prediction of disease course in depression has proven to be a difficult task, with a recent systematic review and meta-analysis showing an average accuracy of 60% for predicting remission or resistance after treatment in adequate-quality studies (included studies used a follow-up time span of 8 to 24 weeks) (11).

One challenge with predicting MDD outcomes is that its etiology and phenotype differ widely between individuals, and large interindividual variation may exist with regard to relevant predictors (12–15). Especially when only a limited set of predictor variables are included for prediction modeling, chances of accurately capturing complex multimodal system dynamics (i.e. the biopsychosocial model of depression) with those variables further decrease. With the availability of novel machine learning methods that can learn complex high-dimensional non-linear patterns in data, a solution to this problem might be to incorporate multiple high-dimensional data sources, each containing putative predictive factors.

While several studies have tried to predict MDD course from a range of different data modalities (e.g. clinical variables, metabolomics, imaging data, epigenetics) (16–20), combining multiple data modalities for predicting MDD chronicity has been relatively scarce. In one recent analysis of the Netherlands Study of Depression and Anxiety (NESDA) cohort (21) that integrated clinical, psychological and biomarker data, predictions of two-year chronicity in MDD reached a balanced accuracy of 62% using a penalized linear model (22). Adding limited biological data showed no improvement in prediction accuracy over the combination of clinical and psychological data (22).

Another NESDA analysis using a similar model with epigenetic data showed an AUC (Area Under the Curve) of 0.571 for predicting the same two-year chronicity outcome (20). Remission after 6 years was predicted more accurately, but the reported AUC (0.724) was based on 10-fold cross-validation results, not on outheld test set results, possibly leading to overoptimistic performance metrics (23,24). Interestingly, neither adding genome wide SNP (single nucleotide polymorphism) data, nor adding 27 clinical, demographical and lifestyle variables improved predictions (20). This is an important finding as no other studies have integrated features from multiple high-dimensional biological data (i.e., multiomic data) and clinical data to improve predictions of MDD disease course. This contrasts with other fields of medicine, where multimodal data integration has led to significant advancements in the field of precision medicine (25), most notably in the field of precision oncology (26,27).

To further investigate the potential of multimodal data in the field of precision psychiatry, the current study explores the potential of integrating multi-omic, clinical, psychological and demographical data. To this end, we used high-dimensional multimodal data collected in 804 NESDA subjects with MDD (21). In a subset of 643 individuals (80% of total sample), using combinations of lipid-metabolomic, proteomic, transcriptomic, genetic, demographic, psychological and clinical data measured at baseline (i.e. from the moment of MDD diagnosis), we used cross-validated machine learning models to predict MDD remission after two years of follow-up. To allow for non-linear pattern detection, and to assess the potential benefit of non-linear models over linear models in multimodal pattern detection, we employed several linear and non-linear machine learning algorithms (elastic net, support vector machine, random forest, XGBoost, artificial neural network). The validity of our models’ predictions was then tested in a separate outheld test group of 161 individuals (20% of total sample).

To embed our machine learning models’ performance metrics in the context of clinical expertise (i.e. how good or bad are predictive performances from a clinicians’ point of view), we additionally let four clinical psychiatrists predict 2-year chronicity in a subset of 200 individuals on the basis of extensive clinical information.

## Methods and Materials

### Participants

In the current study we included data that was collected as part of a larger, multi-center longitudinal study (NESDA, n = 2981, see Supplementary Methods) (21). We included a subsample from the NESDA cohort consisting of 804 subjects with the following inclusion criteria (identical to our previous study (22)): (i) presence of a DSM-IV MDD or dysthymia diagnosis (or both) in the past 6 months at baseline, established using the structured Composite International Diagnostic Interview (CIDI, version 2.1) (28), (ii) confirmation of depressive symptoms in the month prior to baseline either by the CIDI or the Life Chart Interview (LCI) (29); and (iii) availability of 2-year follow-up data on DSM-IV diagnosis and depressive symptoms measures with the CIDI. The ethical review boards approved the NESDA research protocol and all participants signed written informed consent.

We defined two outcome groups: remission or no remission two years after follow-up. We based the outcome on the presence or absence of a current unipolar depression diagnosis (6-month recency MDD diagnosis or dysthymic disorder) at 2-year follow-up, according to DSM-IV criteria. Table 1 lists sample characteristics and statistics for both outcome groups of all 804 included subjects.

**Table 1.**
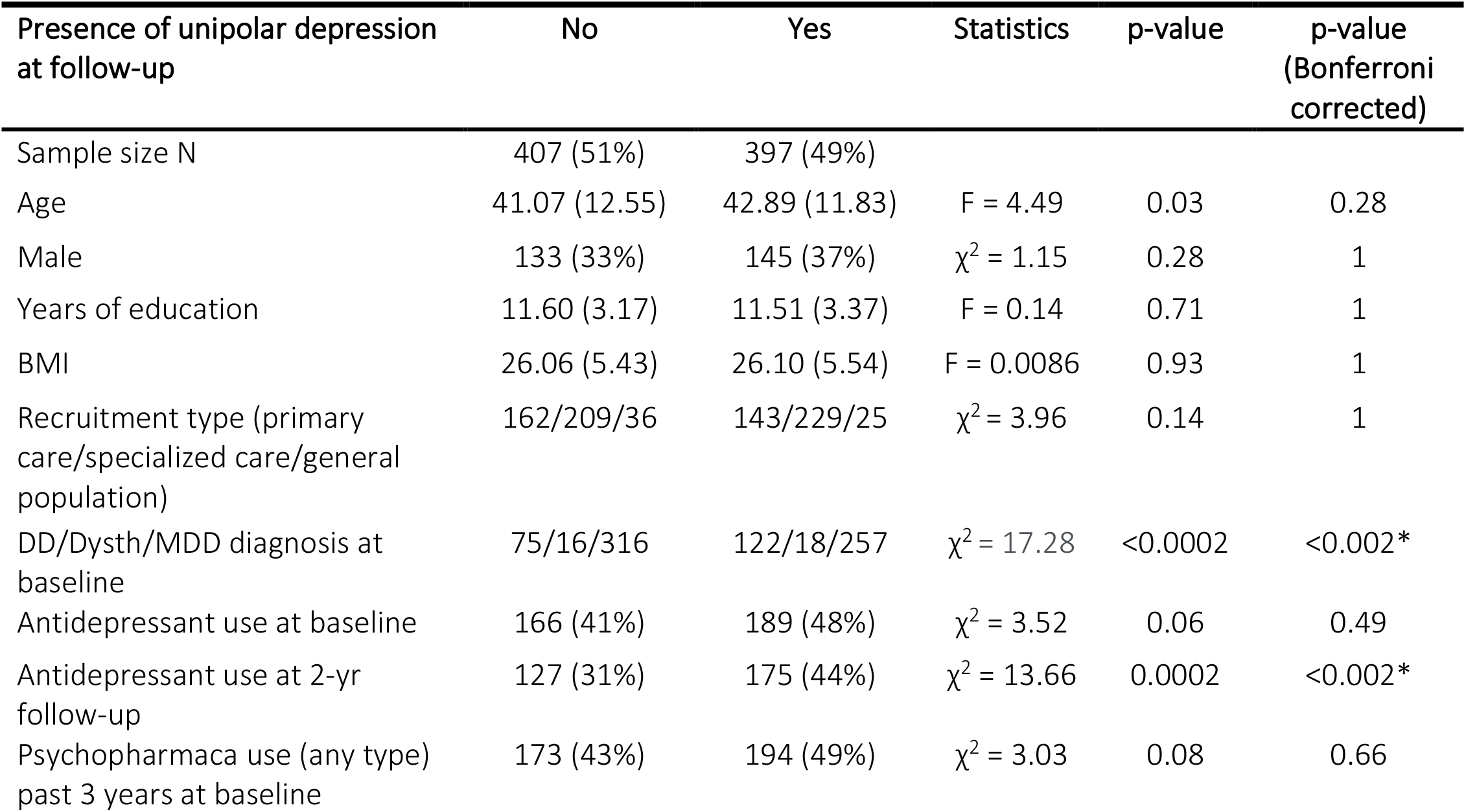
Sample characteristics. The table shows characteristics of the total sample divided by the presence or absence of a unipolar depression diagnosis (major depressive disorder or dysthymia) 2 years after baseline measurement. Data are given as mean (standard deviation) or N (%). *MDD*: major depressive disorder; *Dysth*: dysthymia; *DD* double depression (both MDD and Dysth diagnosis).

### Clinical variables

We included a set of 10 relevant clinical, psychological and demographical predictor variables (to which we will now refer to as ‘clinical variables’), including age, sex, years of education, depressive symptom severity (IDS-SR questionnaire) (30) (both as total score and severity category ranging from one to five) and five personality dimensions (neuroticism, extraversion, openness to experience, agreeableness and conscientiousness), measured with the NEO five-factory inventory (31). Selection of the included clinical variables was based on the results of our previous study in the same NESDA subsample where depressive symptom severity and personality dimensions proved to be most predictive (22).

### Proteomic variables

For 611 of the included 804 subjects, a panel of 243 analytes involved in endocrinological, immunological, metabolic and neurotrophic pathways was assessed in serum at baseline. A full list of the 243 analytes and their inclusion in predictive modelling with missing percentages per variable can be found in Table S1. The Supplementary Methods provides further details on data collection and data processing.

### Lipid and metabolite variables

A lipid-focused metabolomics platform was used to measure 231 lipids, metabolites and metabolite ratios in plasma at baseline for 790 of the included 804 subjects.

From now on, we refer to this data as ‘lipidomic data’. The Supplementary Methods describes further lipidomic data details.

### Transcriptomic variables

Transcriptome-wide expression levels were measured in whole blood for 669 of the 804 included individuals. For each subject, 44 241 microarray probes targeting 23 588 genes were available for analysis. See Supplementary Methods for details.

### Genotype data

Using the LDpred package in R (32), a total of 29 polygenic risk scores (PRSs) were calculated for 701 of the 804 included subjects with available genotype data. Further details about DNA extraction and PRS calculation can be found in the Supplementary Methods. Table S3 lists all 29 phenotypes for which polygenic scores were calculated (e.g. MDD, anxiety, neuroticism etc.).

### Analysis

All analyses were performed using the programming languages R (version 4.0.3) and Python (version 3.8.5). All R and Python code is made publicly available on GitHub at https://github.com/pchabets/chronicity-prediction-depression.

### Machine learning analysis

Full details about data preprocessing, train and test procedures and model evaluation are described in the Supplementary Methods section. In short, first, XGBoost models (33) were trained using each data modality separately to predict 2-year chronicity. Second, to investigate possible prediction augmentation effects of combining clinical and high-dimensional biological data, separate XGBoost models were trained using the combination of clinical data added to each of the separate - omics data sources to predict 2-year chronicity. Third, another XGBoost model was trained using the combination of all data modalities together.

Based on the combination of data that proved to result in the best XGBoost model predictions (i.e. proteomics + clinical data, see *Results* section), we additionally investigated the nature of the multimodal predictive signature by running different linear and non-linear algorithms including elastic net, support vector machine (SVM), random forest (RF) and a feed-forward densely connected artificial neural network (ANN) using 1) only clinical features (i.e. severity scores, psychological and demographical variables); 2) only proteomics data, and 3) the combination of both data modalities. For each model prediction augmentation by adding proteomics data to the clinical data was evaluated using the area under the receiver operating characteristic (AUROC). All performance metrics reported are from validating the trained models on outheld test data (see Supplementary Methods for details).

### Feature importance analysis

Feature importance analysis was based on computing Shapley values for every feature included in the best performing XGBoost model, using the SHAP implementation for XGBoost (34,35). SHAP allows for calculating feature importance levels per individual prediction, and deriving global feature rankings according to those levels (35). Ranking features according to their overall importance is done by using the mean of absolute Shapley values computed for all individual predictions (35). Feature ranking and relation to measured variable values was plotted using the SHAP and SHAPforxgboost packages (35). Protein-protein interaction and enrichment analysis was performed using the metascape platform (36). Further details are provided in the Supplementary Methods section.

### Human predictions

Four human raters (trained and board-certified psychiatrists) independently predicted 2-year chronicity for 200 MDD subjects using clinical baseline data. Each rater was given two sets of samples for prediction. In the first sample, raters had to predict 2-year chronicity status of subjects on the basis of the same 10 clinical baseline predictor variables used by the machine learning models. In the second sample, raters additionally had access to baseline data on 1) dysthymia diagnosis; 2) MDD history; 3) anxiety diagnosis (lifetime); 4) one-month recency of anxiety disorder symptoms; 5) alcohol diagnosis status (lifetime); 6) recency of alcohol abuse or dependency; 7) total disease history (totaling 17 baseline predictor variables).

Further details about the human prediction process and interrater agreement analysis are described in the Supplementary Methods section.

## Results

### Proteomic data is most informative for predicting 2-year chronicity

We first tested how well 2-year chronicity in MDD can be predicted for each data modality separately. For each of the available data modalities, the train, validation and test sets used for the classification models approximated balanced distributions of the two outcome classes (Table S4). For unimodal data predictions, the model using proteomic data showed highest performance (AUROC = 0.67, balanced accuracy = 0.68), followed by the models informed by clinical data (AUROC = 0.63, balanced accuracy = 0.62) and genetic data (AUROC = 0.61, balanced accuracy = 0.60) (Figure 1, Table S4). All models reached accuracy levels significantly above chance level. For the model informed by PRSs, accuracy only reached a significant accuracy level when using a cutoff on the ROC curve that made the model significantly biased towards false negative classifications (McNemar’s test, p = 1.86e-06) (Table S4).

**Figure 1.**
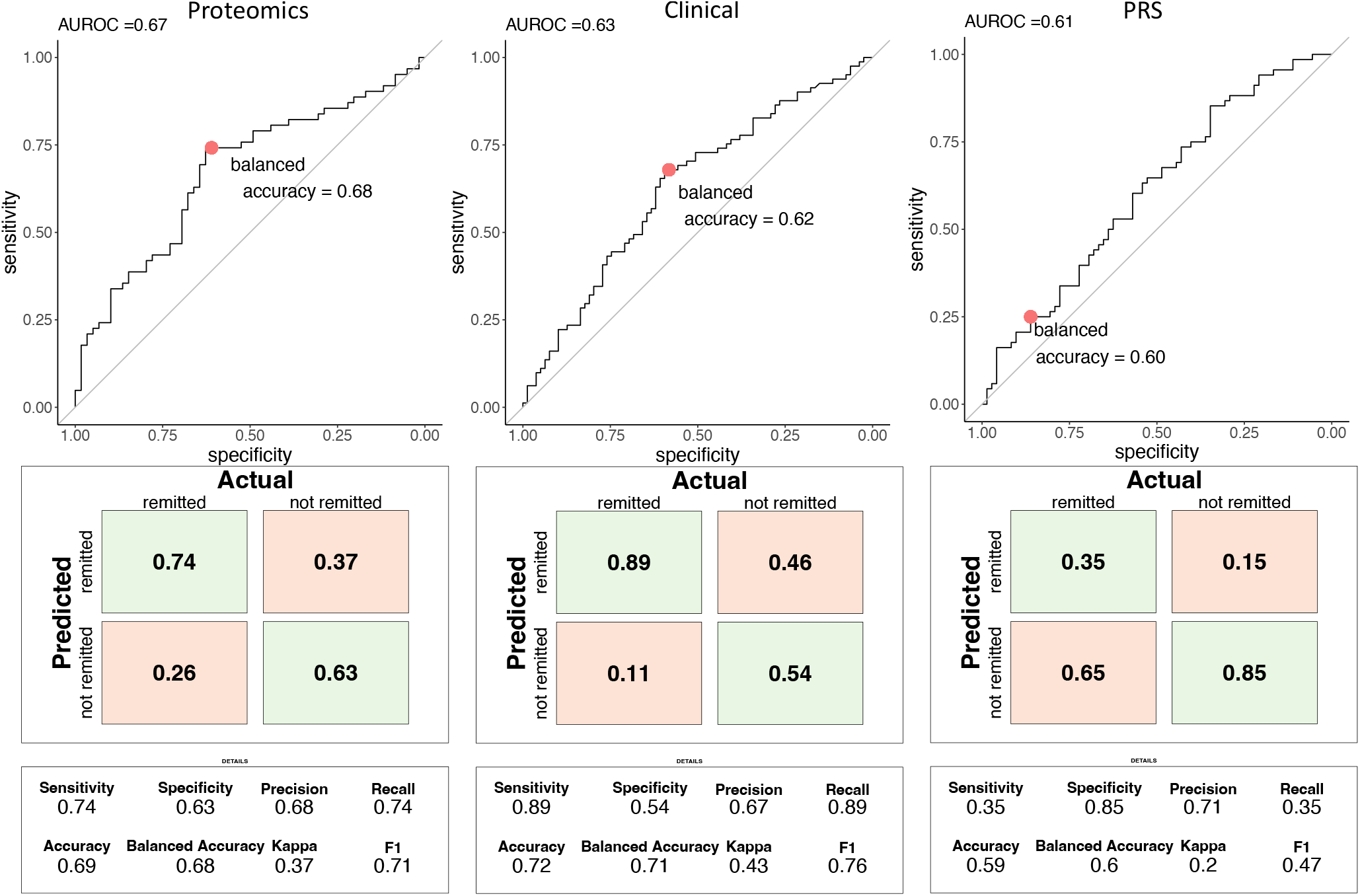
Predictive performance of XGBoost models informed by either proteomic data (left), clinical data (middle), or PRS (genetic) data (right). ROC curves are plotted separately, with the reported area under the ROC curve (AUROC) and the maximum balanced accuracy shown for the optimal class probability cutoff. For each model a confusion matrix is shown with additional performance metrics.

### Combining clinical and proteomic data augments prediction performance

Models informed by both clinical and omics data outperformed models informed by unimodal omics data in every case, most robustly for combining proteomic and clinical data (figure 2). All combinations of clinical and omics data resulted in higher predictive performance than the model informed by clinical data only, except for combining clinical and transcriptomic data (Figure 2). Although a clear trend in augmented predictions by combining omics with clinical data was observed for all omics data (Figure 2), only the augmented performance of adding proteomic to clinical data reached statistical significance (AUROC = 0.78 versus AUROC = 0.63, p = 0.013).

**Figure 2.**
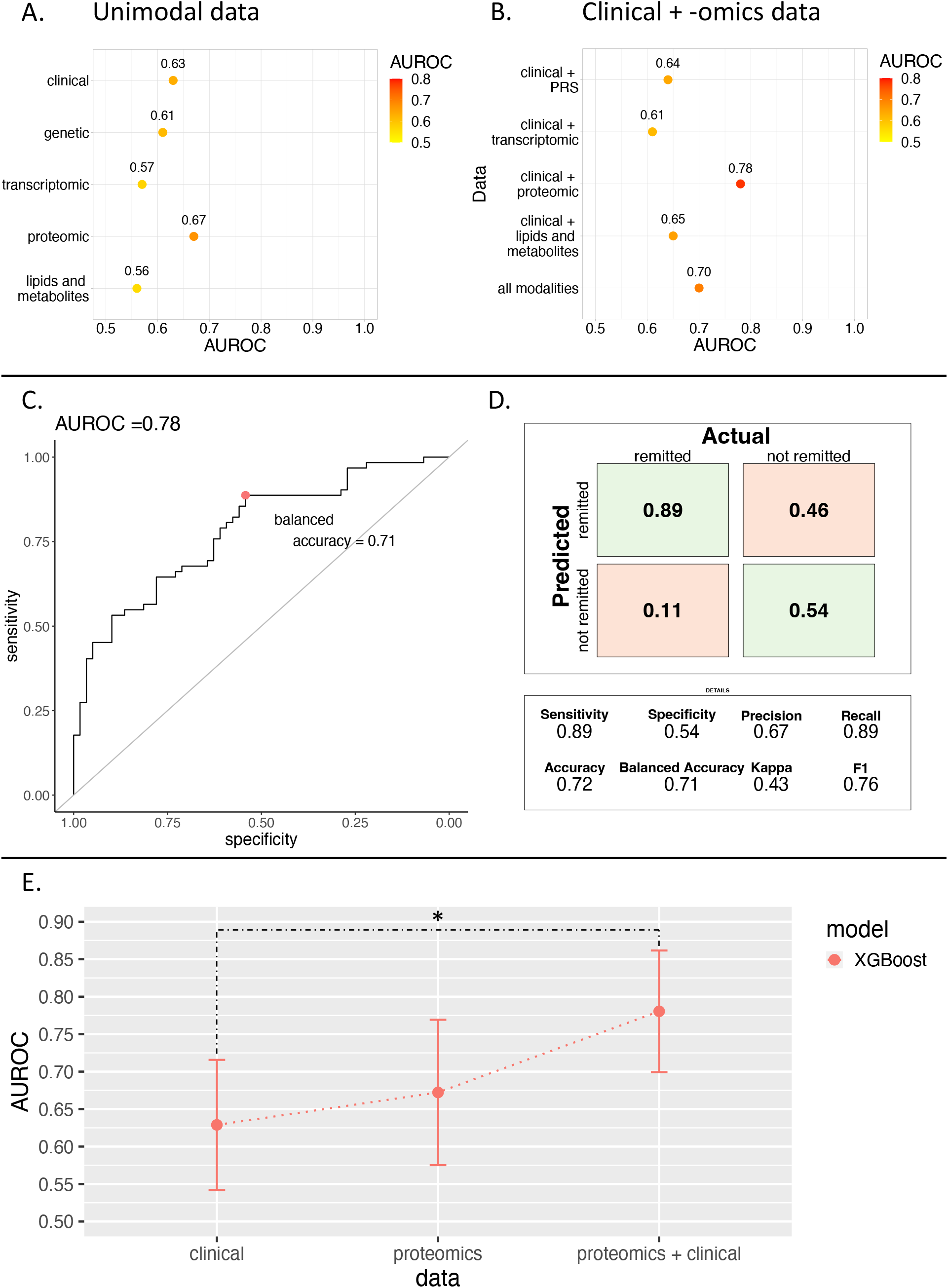
A – Predictive performance of XGBoost models trained on several single data modalities. The y-axis shows the data modality used for training and testing the model. The x-axis shows the AUROC score of each model tested on the outheld test set. **B** – The same as A, but results for multimodal data are shown. **C-D** – ROC-curve plotted with AUROC for the XGBoost model informed by both clinical and proteomic data. The maximum balanced accuracy is shown on the ROC curve, showing the optimal probability cutoff for highest classification accuracy. On the right, the confusion matrix with additional performance metrics is shown for this model. **E** – Performance metrics (AUROC) of the XGBoost models informed by only clinical, only proteomic, or the combination of both are shown, with the 95% confidence interval of the AUROC indicated by the whiskers. A significant difference in AUROC values between model’s is indicated by an asterisk.

To further investigate the augmented prediction of MDD 2-year chronicity when adding proteomic to clinical data, we used several linear and non-linear machine learning models informed by clinical, proteomic, and the combination of both data (Table S4). Informing machine learning models by only proteomic data resulted in low predictive performance for linear models, compared to non-linear models (Figure S2). Augmented predictive performance by adding proteomic data to clinical data was not found for any linear model, but was observed for all non-linear models, most pronounced for XGBoost, and reached statistical significance only for the XGBoost model (p = 0.013, Figure S2).

### Variable importance analysis shows predictive pattern enrichment of analytes involved in inflammatory response and lipid metabolism

SHAP-analysis was performed on the best performing unimodal and multimodal informed models (i.e. XGBoost informed by proteomic data and XGBoost informed by clinical and proteomic data). Both for the proteomics-only model, and the model informed by both clinical and proteomic data, blood fibrinogen levels showed highest mean absolute SHAP values (Figure 3, Table S5). Symptom severity at baseline showed to be the most predictive clinical feature for 2-year chronicity in MDD (Figure 3, Table S5). For the proteomics-only model, 109 analytes had an average absolute SHAP value > 0 (i.e. were informative for predictions). For the combined data model, 42 features were informative for predicting 2-year chronicity in MDD model, including 38 proteomic analytes. Age, years of education and sex were not attributed any SHAP values in the multimodal XGBoost model (Table S5).

**Figure 3.**
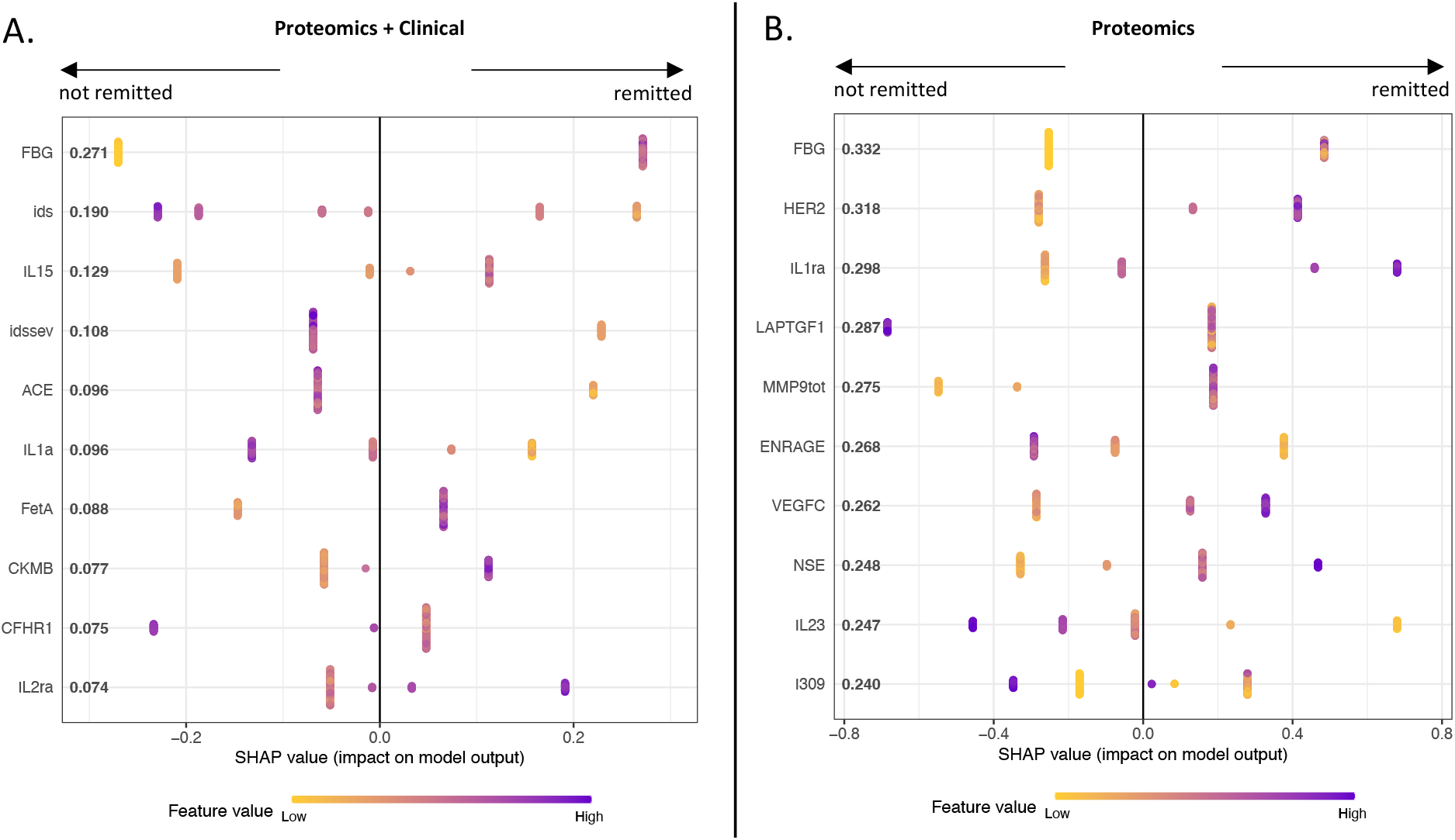
**A** – SHAP analysis results visualized for the XGBoost model informed by both clinical and proteomic data. On the y-axis, the top 10 most important features (ranked by absolute average SHAP value, i.e. global SHAP value) are shown with their respective global SHAP value. Each dot in the graph indicates a single prediction (i.e. one subject). The position of the dots on the x-axis shows the impact that the feature value had for that individual prediction, with a negative SHAP value meaning that the model’s decision was pushed towards a ‘not remitted’ classification, and a positive SHAP value meaning that the feature’s value pushed the model’s decision towards predicting the ‘remitted’ class. Colors indicate the relative feature value measured in a subject (relative to the mean of all subjects), with yellow indicating a relatively low value, and purple a relatively high value. Note that most dots are stacked on each other due to the fact that a range of feature values for several individuals can result in similar SHAP values (i.e., different feature values can influence the model’s decision by the same magnitude and direction). **B** – Similar to A, but here SHAP results of the XGBoost model informed by proteomic data only is shown. Proteomic abbreviations shown on the y-axis are listed by full name in Supplementary Table 1. *IDS: Symptom Severity (continuous measure); IDSSEV: Symptom Severity (categorical measure)*.

Proteomic analytes that were informative in the combined data model and in the proteomics-only model were analyzed separately for protein-protein interactions and pathway enrichments. Network analysis of protein-protein interactions revealed densely connected subnetworks associated with inflammatory response and lipid metabolism for both the unimodal and multimodal XGBoost model, with enrichment of Reactome -, GO - and WikiPathway terms related to interleukin-10 signaling, chemokine signaling pathway, cholesterol esterification and reverse cholesterol transport (see Figure 4).

**Figure 4.**
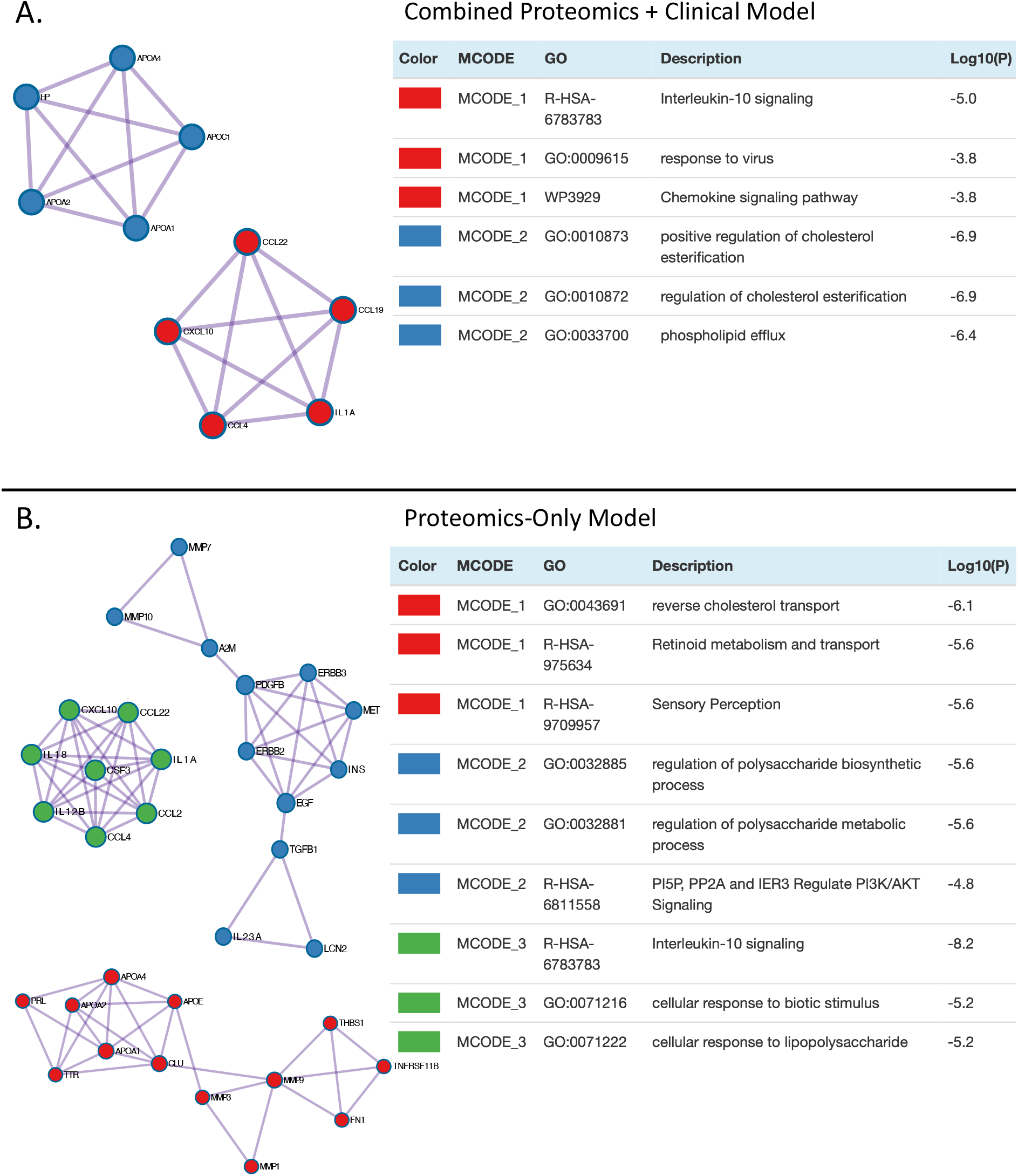
**A** – densely connected protein-protein subnetworks and subsequent enrichment analysis results for the informative proteomic analytes in the combined clinical and proteomic data model. **B** – similar to A, but results of the informative analytes of the proteomics-only model are shown. *GO: gene ontology database; WP: WikiPathways database; RHS-A: Reactome pathway database*.

### Human prediction of 2-year chronicity from clinical data

Four clinical psychiatrists independently predicted 2-year chronicity retrospectively from baseline data for 200 subjects with balanced subjects’ outcome distribution (2-year chronicity n = 100, remitted n = 100). Using the 10 clinical features that the clinical XGBoost model was informed by, human raters had an average accuracy of 0.51 (min = 0.35, max = 0.63, Figure 5). When additional relevant clinical baseline data was available to the human raters, the raters’ average prediction accuracy increased to 0.55 (min = 0.33, max = 0.65, Figure 5). Interrater reliability between the four raters was low (Fleiss’ kappa = 0.32, p = 7.26e-7). Both the XGBoost model trained on the same clinical data, and on combined clinical and proteomics data, outperformed all human predictions (Figure 5).

**Figure 5.**
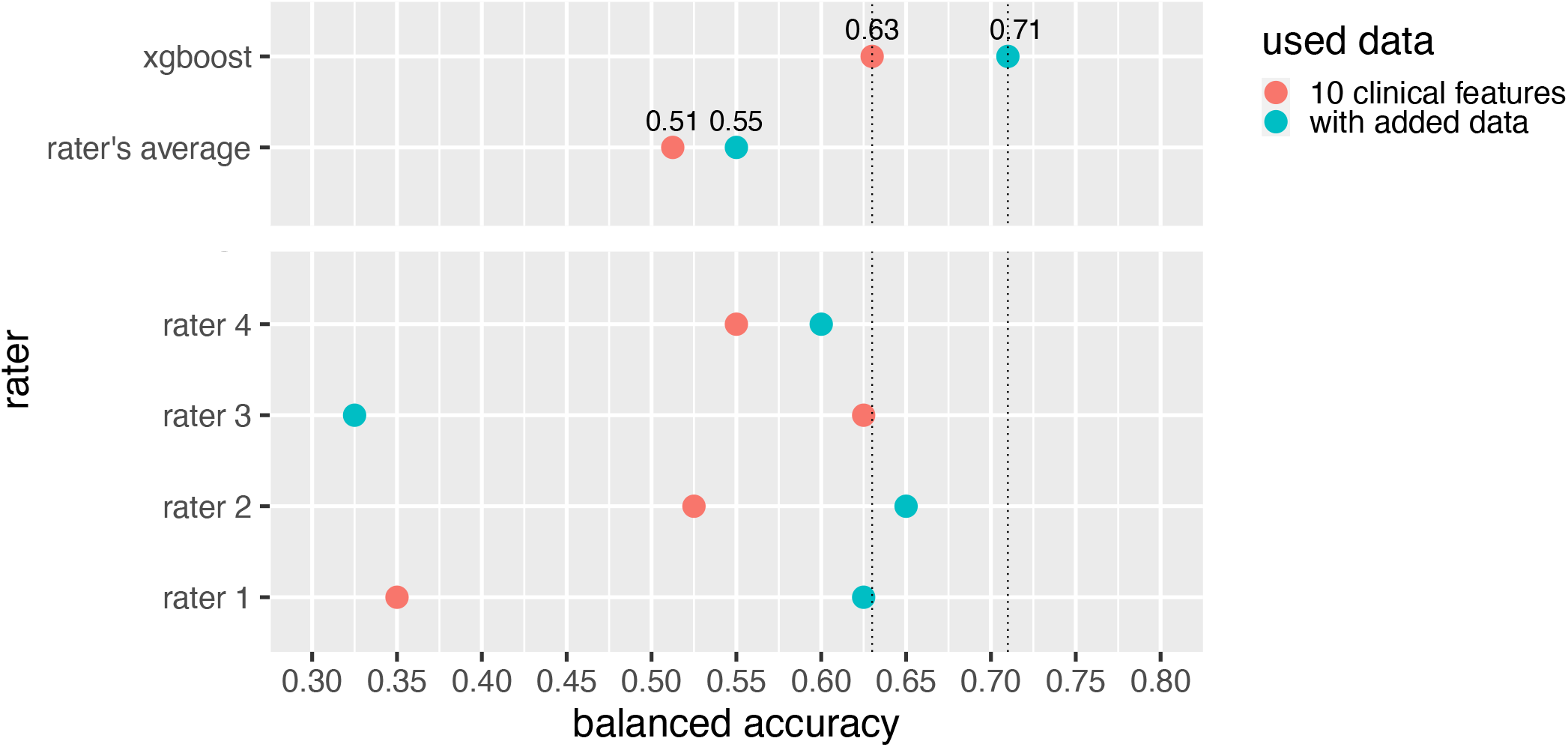
Results of clinical psychiatrist’s predictions of 2-year MDD chronicity, compared to the XGBoost model’s performance. The x-axis shows the balanced accuracy of predictions. The dots represent a psychiatrist’s or model’s predictive performance, with the color indicating on the basis of what information predictions were based.

## DISCUSSION

In this study we showed that longitudinal prediction of 2-year MDD chronicity substantially benefits from integrating multimodal data compared to relying on unimodal data. Specifically, model predictions significantly improved when combining proteomic and clinical data (Figure 2). Our model informed by only clinical data showed identical performance to the previously reported performance of a linear model using multiple data modalities, including those 10 clinical variables, in the same dataset (22). Performance of our model predictions significantly increased when adding proteomic data, but only for non-linear models, suggesting superior multimodal predictive pattern detection by non-linear models over linear models (Figure S2). Subsequent SHAP analysis revealed a multimodal predictive signature consisting of baseline symptom severity, personality traits and peripheral blood biomarkers related to the immune system and lipid metabolism.

### Interpretation of predictive features

Findings of our SHAP analysis are consistent with a previous NESDA study that showed symptom severity to be predictive of MDD chronicity (22). Interestingly, while our proteomic results indicate a predictive inflammatory component, the same study previously found that including three inflammatory markers (CRP, IL6, TNF-alpha) did not improve predictions, indicating the need for higher proteomic resolution (22). Detection of a predictive inflammatory signature is particularly insightful given the established association between low-grade inflammation and MDD (37), and other studies reporting MDD phenotypes associated with inflammatory and metabolomic markers (4,38). Higher levels of fibrinogen (an acute-phase protein) have been related to MDD (39), antidepressant intake (39,40), drug treatment response (41) and remitted MDD (42,43). SHAP analysis of our most predictive models now reveals that fibrinogen levels are part of a multimodal signature predictive of 2-year chronicity in depression, with a general tendency to push the model’s decision closer to the ‘not-remitted’ class when fibrinogen levels are lower (Figure 3). Yet, however tempting, given the nonlinear multimodal nature of the predictive signature, no valid conclusion can be drawn as to whether these results indicate high or low fibrinogen as a risk factor for 2-year chronicity (44). Moreover, considering a group of interacting analytes rather than one isolated analyte can improve biological-mechanistic insights. For this reason, we used protein-protein interaction networks and enrichment analysis (Figure 4). In this light it also deserves mentioning that the other lipid-metabolic and immunomarkers together have considerably higher predictive value than fibrinogen levels alone (e.g. in the combined model, the added absolute mean SHAP values of predictive proteomic analytes excluding fibrinogen equals 1.75, versus 0.27 of fibrinogen alone). It is only the combination of variables that ultimately enables the model to predict with good performance (AUROC = 0.78) – illustrative for the idea that different data modalities provide complementary information (25).

Although a signature of lipid metabolite levels has been associated with MDD (45), and a lipid-related signature was part of the multimodal signature found most predictive of 2-year chronicity, lipidomics as unimodal data source did not result in good 2-year chronicity predictions in MDD. Adding lipidomic measurements to clinical data did result in slight improvement in predictions, although not significant.

These results could indicate that the coverage of our lipidomics platform was not broad enough, or that other metabolites than lipids might be more relevant for prediction. Likewise, one transcriptomic study has shown differential gene expression associations with MDD on a group level (46), but our study showed that these findings did not translate to good individual predictions of 2-year chronicity in MDD. Moreover, adding transcriptomic to clinical data seemed to only add noise to the model, i.e. worsen model prediction performance, although this difference did not reach statistical significance. Although the model informed by all data modalities performed better than the clinical-only model (AUROC = 0.70 vs 0.63), integrating all -omics data with clinical data did not yield improved predictions over the combined proteomics and clinical model, indicating redundancy of the other -omic data in our NESDA sample considering 2-year MDD remission predictions (Figure 2A-B).

Combining several PRSs resulted in prediction accuracy significantly above the chance level, even more so than lipidomic and transcriptomic data, and showed similar performance metrics to the model informed by clinical data only. However, adding this genetic data to clinical data did not substantially augment prediction performance (AUROC = 0.63 vs 0.64, p > 0.3). Although our results did not show any predictive benefit of combining PRS and clinical data, future PRS improvements, combined with more suited genetic dimensionality reduction techniques for machine learning based predictions, could result in the further improvement of multimodal predictions for complex traits.

The observation that the model informed by all data modalities did underperform compared to the model informed by clinical and proteomic data (AUROC = 0.70 vs 0.78) – even though this observed difference was not significant (p = 0.234) – can be explained by 1) the fact that sample size for the multimodal informed model was considerably less whilst including more features (features = 254 vs features = 192), meaning that the model using all data modalities ideally needed more, but learned from less samples during training (n = 405 vs n = 490), and 2) because the inclusion of data modalities that were less predictive than the combination of clinical and proteomic data likely added noise to the model, resulting in suboptimal results.

Ultimately, the only model significantly outperforming the model based on clinical data was the model informed by both clinical and proteomic data. Additionally, proteomics data was most informative for unimodal data predictions. Although proteomic data had a larger feature space than the clinical and PRS data, this was not the case compared to the lipidomic and transcriptomic data that showed to be less predictive. Consequently, the superiority of proteomics-informed predictions in our study cannot solely be explained by dimensionality. Illustratively, the previous NESDA analysis predicting 2-year chronicity in the same set of subjects used a substantial larger amount of clinical variables (55 variables), but showed no improvement in predictions compared to our model based on only 10 clinical variables (balanced accuracy 62% for both models) (22). Indeed, dimensionality does not equal informativeness. Provided that a recent NESDA analysis including high dimensional epigenetic data showed poor 2-year chronicity predictions (AUROC = 0.571) (20), our current study indicates that, considering multiple -omic data modalities, proteomic biomarkers might be most informative for MDD 2-year chronicity prediction – most notably when combined with clinical data. Given the currently limited sample sizes and robustness of neuroimaging studies predicting clinically relevant MDD outcomes (47–49), and the difficulty of integrating imaging as a routine clinical practice, our results warrant consideration of (immune and lipid metabolism focused) proteomics as a feasible approach to chronicity predictions that show potential for clinical relevance.

### Prediction and models’ performance

Applying machine learning models entails predicting at the individual subject level (n=1 predictions), which might ultimately pave the way for individual clinical application, i.e. enable personalized psychiatry (50). For personalized psychiatry, accuracy and other prediction performance metrics are arguably more valuable than traditional statistical measures because they 1) indicate how well a model works *on the individual subject level* and 2) are the result of the model being *put to practice* in separate ‘new’ individuals, not previously ‘seen’ by the model.

One might regard a balanced accuracy of 71% (i.e. our best performing model) as too low for use in clinical practice. In line with previous findings (51), we however showed that the next best thing for patients – namely interpretive predictions by clinicians – did perform substantially worse. Moreover, we showed clinician’s predictions to show high interrater variability, resulting in low interrater reliability (kappa = 0.32) (52). One can argue that the retrospective data shown to the clinicians in our study does not approximate a live clinical impression. Previous studies have however shown that this added source of information for predictions results in even worse predictions by clinicians (53,54). This does not mean live clinical impressions are uninformative for future predictions per se. In the light of multimodal prediction, live clinical impressions might yet prove to hold complementary information for augmented predictions. Future studies will have to clarify for what type of outcome predictions (e.g. therapy response, remission), and in combination with what type of data, clinical impressions add a complementary layer of predictive information.

A next much needed step for personalized psychiatry is to implement and test machine learning based clinical decision making in clinical trials. Clinical implementation of machine learning models showed promise in preliminary studies (55,56). Importantly, recent clinical trials successfully showed the superiority of machine learning-based clinical decision making compared to conventional clinical decision making for medical fields outside of psychiatry (57,58). To facilitate in future prospects of personalized, machine learning-aided decision making in psychiatry, we view our study as an important clue for the type of data that can be informative for prediction models put into practice, specifically with the aim of predicting an individual’s naturalistic course of MDD from baseline data. Such predictions may ultimately benefit patients’ outcomes by providing clinically actionable information.

For example, in cases predicted to show a chronic disease progression, intensifying therapy early on might improve disease course.

### Strengths and limitations

We were able to train, (cross-)validate and test our models on subject data collected as part of the longitudinal NESDA study. Likewise, human prediction evaluation was solely based on data of subjects included in the NESDA database. Unfortunately, without any validation of our model on data external to the NESDA dataset, robust assessment of the generalizability of our model’s performance is currently lacking.

Given that we carefully prevented any data leakage from final test set to the train set in all our imputation, preprocessing and feature selection procedures (i.e. prevented ‘double dipping’) (23,59,60), used balanced train and test sets (61), used separate repeated 10-fold cross-validation procedures in the train sample independent of the final test set (23,24), used a sample size for prediction analysis of several hundreds (23), and tested final model performance on an outheld test sample (23,47,60), we however believe that, at the least, the multimodal features that were found to be most predictive represent robust findings.

We therefore argue that the importance of this work primarily lies in the observations that 1) there are blood-based variables that can be individually predictive for the naturalistic course of MDD; 2) individual predictions become more accurate when using multi-modal data; 3) high-dimensional predictive signatures might not be detected using conventional linear machine learning models. Secondary, zooming in on the multimodal predictive signature that resulted in the best predictions, this signature was found to consist of baseline symptom severity, personality traits and peripheral blood biomarkers related to the immune system and lipid metabolism. These results are useful in guiding variable inclusion decisions in future MDD chronicity prediction studies.

We chose 2-year chronicity instead of a longer period of enduring depression because i) Persistent Depressive Disorder is defined as depression lasting for at least 2 years in DSM-5 300.4 (F34.1) (62), and ii) sufficient sample size was available only for the 2-year time period (although still suboptimal for the model informed by all modalities). Inherently, our ‘chronicity’ label is based on a somewhat arbitrary cutoff, both by time span and diagnostic criteria, that might condense truly chronic phenotypes from delayed remission cases in the same group. This could be an explanation for prediction performance that is mediocre at best when using unimodal -omics data, and below excellent when using multimodal data.

## Conclusion

To our knowledge, this is the first study to show that the combination of multimodal biological and clinical data significantly improves the accuracy of individual predictions of 2-year chronicity in MDD in a relatively large sample (total n = 804). Moreover, this study shows that what is predictive of remission of MDD within 2 years is a combined signature of symptom severity, personality traits and immune and lipid metabolism related proteins at baseline. We argue that future studies that investigate the potential of clinical application of MDD course prediction models are much needed, and should consider including both clinical and proteomic data focused on immune and lipid metabolism markers in their data.

## Acknowledgments

This work was supported by the Geestkracht program of the Netherlands Organization for Health Research and Development (Zon-Mw, grant number 10-000-1002) and is also supported by participating universities and mental health care organizations (VU University Medical Center, GGZ inGeest, Arkin, Leiden University Medical Center, GGZ Rivierduinen, University Medical Center Groningen, Lentis, GGZ Friesland, GGZ Drenthe, Institute for Quality of Health Care (IQ Healthcare), Netherlands Institute for Health Services Research (NIVEL), and Netherlands Institute of Mental Health and Addiction (Trimbos). The collaboration project is co-funded by the PPP Allowance made available by Health∼Holland, Top Sector Life Sciences & Health, to stimulate public-private partnerships. The authors would like to thank Dr. E Bosdriesz and Dr. F Bennis for their valuable input on the machine learning analysis, and Dr. J Tijdink and Dr. J Luykx for their aid in complementing human predictions.

