## Supplemental Figures for "Multimodal Data Integration Advances Longitudinal Prediction of the Naturalistic Course of Depression and Reveals a Multimodal Signature of Disease Chronicity"

### SUPPLEMENTARY FIGURES

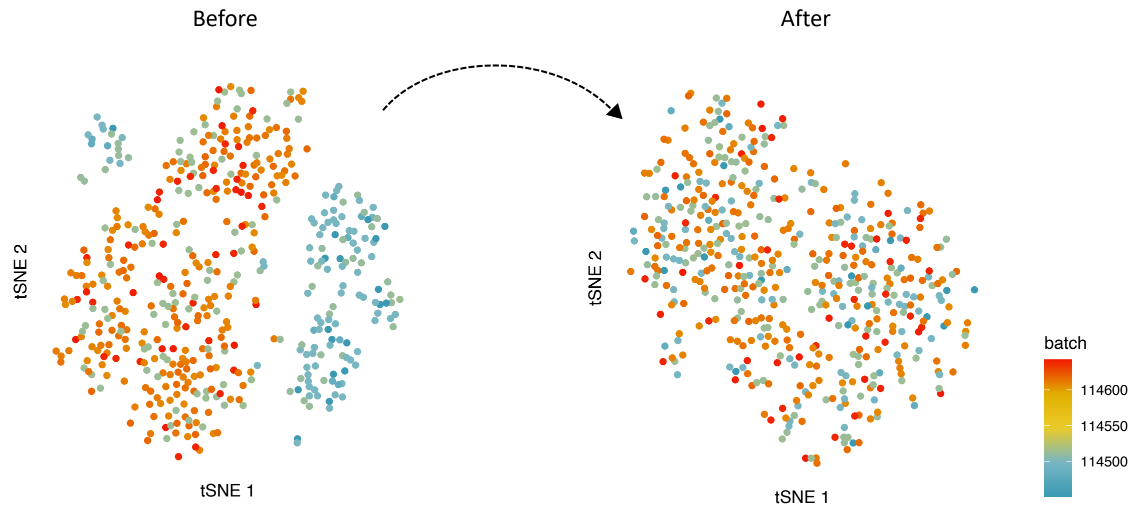


*Supplementary Figure 1* – Results of batch effect removal in proteomics data, visualized as tSNE on the dataset used for training and cross-validation (excluding outheld test set). Before batch correction, tSNE clearly shows separation of samples in subgroups according to batch plate. After batch correction, samples show no clustering by batch.

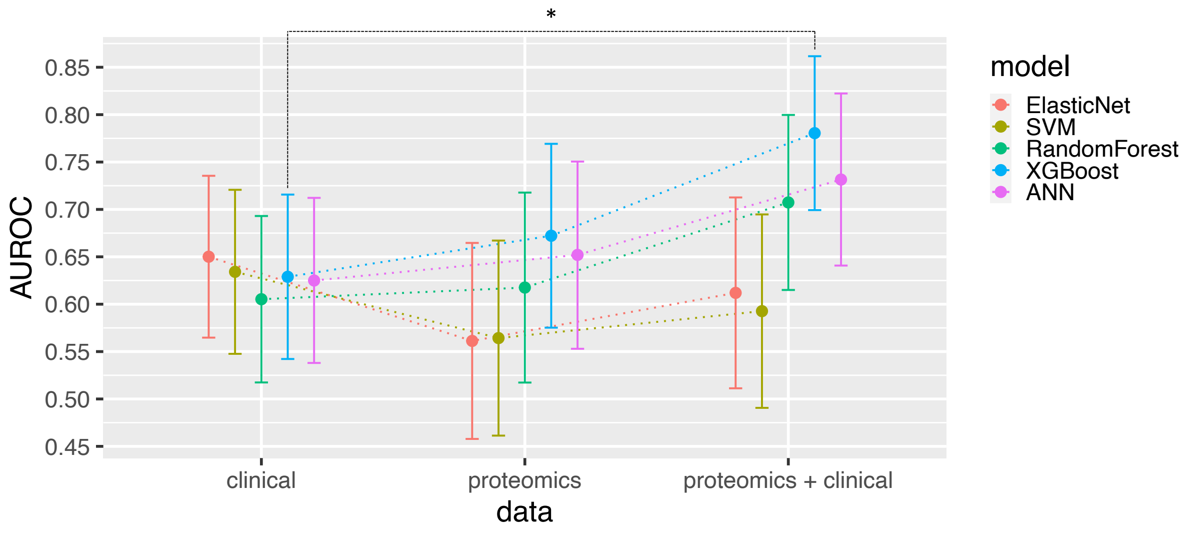


*Supplementary Figure 2* - AUROC values and 95% confidence intervals calculated for several machine learning models informed by either clinical, proteomics, or both types of data. *SVM: support vector machine (linear kernel), ANN: artificial neural network (deep learning model), *: $p < 0.05$*
